# Genetic variation of *Aedes aegypti* populations from Ecuador

**DOI:** 10.1101/2019.12.17.875591

**Authors:** Varsovia Cevallos, Denisse Benítez, Josefina Coloma, Andrés Carrazco, Chunling Wang, Susan Holecheck, Cristina Quiroga, Gabriela Castillo, Britney Tillis, Patricio Ponce

**Affiliations:** Instituto Nacional de Investigación en Salud Pública, Centro de Investigación en Vectores Artrópodos (CIREV), Quito, Ecuador; Universidad Yachay Tech, Escuela de Ciencias Biológicas e Ingeniería. Urcuqui, Ecuador; School of Public Health, Division of Infectious Diseases and Vaccinology, University Berkeley, California, USA; School of Life Sciences, Arizona State University, Tempe, USA; Simon A. Levin Mathematical, Computational and Modeling Sciences Center, Arizona State University, Tempe, USA

## Abstract

This is the first genetic analysis in Ecuador of *Aedes aegypti* using fragments of mitochondrial genes, NADH dehydrogenase subunit 4 (ND4) and cytochrome oxidase subunit I (COI). A total of 154 mosquitoes from 23 localities were collected in the Pacific coastal lowlands, Amazon basin lowlands, and the Galápagos Islands from 2012 to 2019. The analysis of fragments of the genes COI (672 bp) and ND4 (262 bp) and concatenated analysis of both COI and ND4 showed two haplotypes (H1, H2) present in Ecuador mainland and the Galápagos Islands. The phylogenetic analysis identified two well-supported clades. Combined analysis of both genes from ten localities also resulted in two haplotypes. Nucleotide diversity, neutrality tests (Tajima’s test D, Fu and Li’s F*and D*) and AMOVA analysis of the entire data set suggest balancing selection for both genes. The results indicate genetic variation without geographical restriction. COI-H1 grouped with sequences from the Americas, West and Central Africa, East Africa, Asia, and Australia. ND4-H1 grouped with similar sequences from the Americas, Asia and West Africa. COI-H2 grouped with sequences from Asia and the Americas. ND4-H2 grouped with sequences from the Americas. We report overlapping peaks in four sequences that suggest heteroplasmy in the individuals. The origin of the populations of *Aedes aegypti* in Ecuador show African genetic origin and are widely present in several countries in the Americas. One of the genetic variants is more common in all the localities and the two haplotypes are distributed indistinctly in the three geographical sampled areas in Ecuador.

## Introduction

Mosquito-borne diseases represent an important risk to human populations, health systems, and economics, especially in tropical poor countries. *Aedes aegypti* (Linnaeus, 1762) is the main vector of several viruses including Yellow Fever (YF), Dengue (DENV), Chikungunya (CHIKV) and Zika (ZIKV) worldwide. It is estimated that 390 million dengue infections occur each year (Bhatt et al. 2013). For 2018, 561.354 cases of Dengue, 55.329 cases of ZIKV, and 123.087 cases of Chikungunya were reported in the Americas (MSP 2019).

*Aedes aegypti* apparently originated in Africa where it is a complex of species (Tabachnick and Powell 1978) and spread widely to tropical countries during the 17th and 18th centuries. The trade between the Old and New World and the arrival of African people into the Americas contributed to the distribution of new mosquito species into the New World (Halstead 2008). There are two widely recognized subspecies of *Aedes aegypti sensu lato (s.l.)*: the ancestral African type *Ae. aegypti formosus* (Aaf) and *Ae. aegypti aegypti* (Aaa) the worldwide type spread outside Africa, with remarkable epidemiological relevance due to its anthropophilic characteristics (Mattingly 1957, Powell et al. 2018, Tabachnick and Powell 1978).

The epidemiological history of *Ae. aegypti* in the Americas started between 1600 - 1946 with the introduction of a dengue-like disease, followed by a successful regional plan for its eradication mostly with DDT, between 1947 – 1970, that later failed with a massive re-infestation and dengue outbreaks between 1971 and 2010 (Brathwaite-Dick et al. 2012). *Aedes aegypti* was pointed as the vector responsible of the transmission of dengue in the region including Bolivia (1987), Paraguay (1988), Ecuador (1988), and Peru (1990) (Pan American Health Organization 1993, Wilson and Chen 2002). *Aedes aegypti* has been the only known vector of DENV, CHIKV, and ZIKV in Ecuador until recently, when *Aedes albopictus* (Skuse 1894) was also reported (Ponce et al. 2018).

Ecuador has a wide variety of ecosystems in the continental territory and at the Galapagos Islands. The continental territory is divided by the Andes Mountains that extend north to south with three resulting well defined geographic zones: 1) the Pacific coastal lowlands, 2) the Andean highlands, and 3) the Amazon basin. In general, the Pacific coast and the Amazon basin lowlands present climatic, ecological and socio-economic conditions that favor the establishment of *Ae. aegypti* populations (Schafrick et al. 2013), and consequently important outbreaks of Zika, dengue and chikungunya (MSP 2018).

During the last three decades, the genetic identity of *Ae. aegypti* populations has been evaluated in several South American countries using a wide spectrum of genetic markers, which range from allozymes to nuclear DNA and mitochondrial (mtDNA) (Ayres et al. 2004, Costa-da-Silva et al. 2005, Paduan and Ribolla 2008, Powell and Tabachnick 2013).

Mitochondrial genes are widely used for identifying genetic variants, dispersal patterns, phylogeny and population dynamic studies of *Aedes aegypti* (Mousson et al. 2005, Jaimes-Dueñez et al. 2015). Among the genes commonly used are the NADH dehydrogenase subunit 4 (ND4) and the mitochondrial cytochrome oxidase 1 gene (mtCOI). The latter has been widely reported as a useful tool for genetic studies, such as DNA barcoding and identification of mosquito species (Chan et al. 2014), including *Ae. aegypti* in different geographic regions (Calvez et al. 2016, Yohan et al. 2018).

Despite the importance of *Ae. aegypti* as the main vector of DENV, ZIKV, CHIKV and potentially of YFV, there is limited information regarding its genetic variability in Ecuador. The genetic characterization of *Ae. aegypti* natural populations in Ecuador and the description of biogeographical patterns and phylogeographic relationships may contribute to the understanding of the spread of mosquito populations and consequently arboviral diseases transmission.

In this study, we analyzed the genetic diversity of *Ae. aegypti* collected in 23 cities and towns in Ecuador, including the Galapagos Islands, using the cytochrome oxidase subunit I (COI) and the NADH dehydrogenase subunit 4 (ND4).

## Materials and Methods

### Mosquito samples

A total of 154 individuals of *Aedes aegypti* from 22 cities and towns in 14 provinces of Ecuador were collected between 2012 and 2019. Coordinates of each sampling site were registered (Table 1). Mosquito collection included locations in the Pacific coastal lowlands, Amazon basin lowlands, and the Galápagos Islands (Fig. 1). The collecting locations differ in their ecological characteristics and all of them had records of vector-borne viral diseases. Mosquito sampling took place in households in areas considered as high-risk sites for arbovirus transmission due to the high number of arboviral clinical cases reported by sanitary authorities (MSP 2013, 2017). Mosquitoes samples were collected in 14 cities and towns in the coastal Pacific lowlands, six cities in eastern Amazon basin lowlands, and two cities in the Galápagos islands (Fig. 1). Samples were collected as larvae and pupae from peridomestic and domestic breeding artificial containers that were taken to the laboratory for adults to emerge. Adult specimens collected in the field were preserved individually in ethanol (70%) at 4°C.

**Fig. 1.**
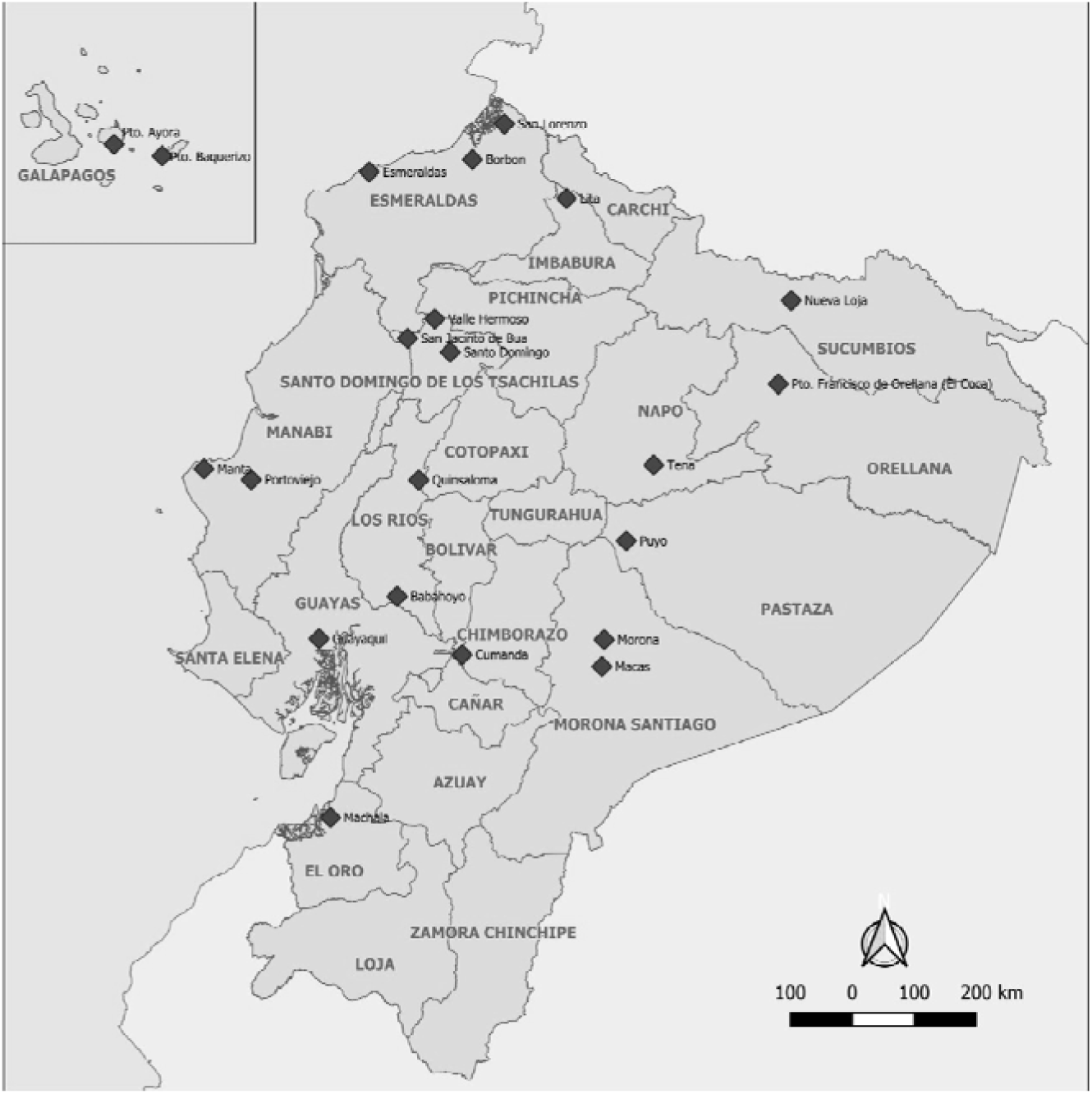
Map of Ecuador showing the localities in the continent and in the Galápagos Islands where *Aedes aegypti* was collected.

**Table 1.** Location, coordinates, date of collection and sample size of Aedes aegypti individuals analyzed for COI and ND4 genes.

| COI | Longitude | Latitude | Date | Sample size |
| --- | --- | --- | --- | --- |
| **Pacific coast** |  |  |  |  |
| Borbón | -78.987 | 1.093 | 2018 | 14 |
| Cumandá | -79.135 | -2.208 | 2014 | 4 |
| Guayaquil | -79.921 | -2.246 | 2016-2017 | 4 |
| Lita | -78.451 | 0.869 | 2018 | 9 |
| Machala | -79.927 | -3.259 | 2017 | 3 |
| Manta | -80.732 | 0.955 | 2017 | 1 |
| Quinsaloma | -79.310 | -1.204 | 2017 | 2 |
| San Jacinto del Bua | -79.518 | -0.101 | 2017 | 1 |
| San Lorenzo | -78.824 | 1.283 | 2017 | 1 |
| Valle Hermoso | -79.278 | -0.085 | 2014 | 1 |
| **Amazon basin** |  |  |  |  |
| Francisco de Orellana | -76.986 | -0.473 | 2015 | 1 |
| Morona | -78.133 | -2.323 | 2015 | 3 |
| Nueva Loja | -76.877 | 0.064 | 2017 | 1 |
| Puyo | -77.956 | -1.479 | 2017 | 1 |
| Tena | -77.820 | -0.982 | 2015 | 2 |
| **GALAPAGOS** |  |  |  |  |
| Puerto Ayora | -90.325 | -0.715 | 2014 | 5 |

| ND4 | Longitude | Latitude | Date | Sample size |
| --- | --- | --- | --- | --- |
| **Pacific coast** |  |  |  |  |
| Babahoyo | -79.679 | -1.787 | 2015 | 5 |
| Borbón | -78.987 | 1.093 | 2018 | 10 |
| Cumandá | -79.135 | -2.208 | 2014 | 8 |
| Esmeraldas | -79.660 | 0.947 | 2014 | 5 |
| Guayaquil | -79.921 | -2.246 | 2016-2017 | 39 |
| Lita | -78.451 | 0.869 | 2018 | 8 |
| Machala | -79.927 | -3.259 | 2017 | 7 |
| Manta | -80.732 | 0.955 | 2017 | 4 |
| Portoviejo | -80.461 | -1.080 | 2017 | 1 |
| Quinsaloma | -79.310 | -1.204 | 2017 | 2 |
| Santo Domingo | -79.156 | -0.222 | 2013-2014 | 3 |
| San Lorenzo | -78.824 | 1.283 | 2017 | 1 |
| **Amazon basin** |  |  |  |  |
| Francisco de Orellana | -76.679 | -0.468 | 2013 | 4 |
| Macas | -78.133 | -2.323 | 2013 | 6 |
| Nueva Loja | -76.877 | 0.064 | 2013 | 8 |
| Puyo | -77.956 | -1.479 | 2012-2013-2014 | 3 |
| Tena | -77.820 | -0.982 | 2015 | 5 |
| **GALAPAGOS** |  |  |  |  |
| Puerto Baquerizo Moreno | -89.594 | -0.910 | 2014 | 3 |
| Puerto Ayora | -90.325 | -0.715 | 2014 | 8 |

| COI+ND4 | Longitude | Latitude | Date | Sample size |
| --- | --- | --- | --- | --- |
| **Pacific coast** |  |  |  |  |
| Borbón | -78.987 | 1.093 | 2018 | 9 |
| Cumandá | -79.135 | -2.208 | 2014 | 2 |
| Guayaquil | -79.921 | -2.246 | 2016-2017 | 2 |
| Lita | -78.451 | 0.869 | 2018 | 6 |
| Machala | -79.927 | -3.259 | 2017 | 2 |
| Manta | -80.732 | 0.955 | 2017 | 1 |
| Quinsaloma | -79.310 | -1.204 | 2017 | 2 |
| San Lorenzo | -78.824 | 1.283 | 2017 | 1 |
| **Amazon basin** |  |  |  |  |
| Tena | -77.820 | -0.982 | 2015 | 2 |
| **GALAPAGOS** |  |  |  |  |
| Puerto Ayora | -90.325 | -0.715 | 2014 | 2 |

COI gene sequences were analyzed from mosquitoes from 16 localities, while ND4 gene sequences were obtained from mosquitoes from 19 localities (Table 1). Phylogenetic trees were built with the COI and ND4 gene fragments as well as with the concatenated sequences (COI+ND4) to determine genetic relationships among the obtained *Ae. aegypti* sequences and the regional and global reported sequences.

### DNA extraction and PCR amplification of mitochondrial COI and ND4 genes

#### Mosquito samples and DNA extraction

Whole body mosquitoes were triturated individually using a homogenizer. *Aedes* DNA was extracted using DNeasy Blood & Tissue Kit® (Qiagen), following the manufacturer recommendations for animal tissue. Amplification for the ND4 gene was carried out in 25 µL of a reaction mixture containing buffer 1X, 0.25 mM of each dNTP, 2mM MgCl2, 0.3 µM of each primer (Forward 5□-GTDYATTTATGATTRCCTAA-3□ and reverse 5□-CTTCGDCTTCCWADWCGTTC-3□) [18], 1.5 U/µl of Taq DNA polymerase, and 5 µL of template DNA. The PCR conditions included an initial incubation at 92°C for 3 minutes, 10 cycles of 92°C for 30 seconds, 48°C for 1 minute, and 72°C for 40 seconds, followed by 40 cycles of 92°C for 30 seconds, 52°C for 35 seconds, and 72°C for 40 seconds, and a final extension at 72°C for 5 minutes. The amplified product was a fragment of 389 bp (ND4 whole CDS: complement 8027..9370, NCBI GenBank:EU352212.1).

COI gene was amplified using the protocol of Paupy et al. (2012) with some modifications. The COI-FOR (5’-TGTAATTGTAACAGCTCATGCA-3’) and COI-REV (5’-AATGATCATAGAAGGGCTGGAC-3’) primers were used. PCR was performed in a 50 μL reaction volume containing 10 μL of 1x buffer, 1.5 mM MgCl2, 0.2 mM dNTP, 0.3 μM of each primer, 1 U/uL of Taq polymerase and 2 μL of DNA template. The PCR cycling conditions included an initial denaturation step at 94°C for 2 min, followed by 35 cycles of 94°C for 30 sec, 54°C for 30 sec, 72°C for 1 min and a final extension of 72°C for 5 min. The amplified product was a fragment of 861 bp (COI whole CDS: 1298.2834, NCBI GenBank:EU352212.1).

### Sequence and phylogenetic analysis

PCR products were detected by agarose gel electrophoresis in TAE 1X buffer, stained with SYBR® Safe 10000X in agarose gel, and visualized under UV light. PCR products were purified and sequenced using the Sanger methodology at Macrogen sequencing service, Seoul, South Korea, UC Berkeley Sequencing Facility or Biodesign Institute, CLAS Genomics Core at Arizona State University.

All sequences were cleaned and aligned using the software Geneious 2019.1.1 (https://www.geneious.com). The obtained sequences were analyzed using BLAST on the NCBI website (http://blast.ncbi.nlm.nih.gov/Blast.cgi) to confirm identity with Ae. aegypti which was between 98 and 100% and a query coverage of 100%. For COI analyses 53 sequences of 672 bp and for ND4 gene 132 sequences of 262 bp were selected. *Culex quinquefasciatus* (GenBank accession KJ012173.1 for COI and GU188856.2 for ND4) was considered as an outgroup for the analyses given that it is closely related in the Culicidae family.

Mega 7.021 (Tamura et al. 2007) was used to select the best-fit model of nucleotide substitution for each gene. For COI the Hasegawa-Kishino-Yano (HKY) model with uniform evolutionary rate of base substitutions was selected. HKY model with gamma distributed rate variation among sites was selected, and 5 discrete Gamma categories were used to analyze ND4. For the analysis of combined COI and ND4 genes the HKY model was selected.

### Phylogenetic trees

Bayesian inference of phylogeny analyses were performed using the BEAST v1.8.3 package (Suchard et al. 2018). BEAUTi was initially used to produce a valid configuration file for BEAST with the selected models from MEGA for each case. Haplotypes trees were inferred using a Tree Prior of Coalescent: Constant Size Multispecies Coalescent (BEAST v1. 7.5, Drummond et al. 2012). For these analyses a strict molecular clock was applied and the default Priors were selected. For analyses of each gene, the length of MCMC chains was of 10 000 000 steps with data and trees sampled every 1000 steps. The generated .xml files were then analyzed in BEAST. The .log files were run in Tracer V.1.7.1 in order to evaluate the coherence of the analysis (Rambaut et al., 2018). The trees files were combined in LogCombiner v1.7.5. To summarize the posterior distribution of tree topologies the Maximum clade credibility tree model was selected in TreeAnnotator v1.7.5, 2001 was the specified number of trees for burn-in (Drummond et al. 2012). Finally, the tree was visualized in FigTree v.1.4.4 (http://tree.bio.ed.ac.uk/software/figtree/).

### Genetic Analyses

For testing the hypothesis that all mutations are selectively neutral the Tajima’s D and Fu & Li F* & D* tests were calculated using DNAsp v.6.12.03 (Librado and Rozas 2009). The same program was used for the estimation of parameters of genetic diversity in all the data: haplotype diversity (Hd), nucleotide diversity (π), the proportion of segregating sites (S) and the average number of nucleotide differences (k). These analyses make it possible to assess the polymorphism patterns observed in the 132 sequences (for the ND4 gene) and 53 (for the COI gene). The TCS network inference method in PopART was used for estimating the inter-haplotype relationship (Clement et al. 2000). Genetic population structure was investigated by hierarchical and non-hierarchical AMOVA and pairwise *Fst* statistics, implemented in Arlequin 3.01 (Excoffier et al. 1992). Molecular variation analysis (AMOVA) was applied to all COI and ND4 sequences to measure population differentiation and the genetic variability. According to Excoffier et al. (1992) is a technique that allows to determine the amount of variation due to the population substructure given an a priori set of population hierarchies where the levels of fixation indexes are estimated in three ways: F_ST_ is used to estimate the proportion of genetic variability found among populations. F_SC_ indicates the internal variability of each population within a same group. F_CT_ shows the variability of each group in relation to the total variation (Costa-da-Silva et al. 2005). Two partitions or estimations were run, a non-hierarchical with all the populations, and other among the populations of the Pacific coastal lowlands, Amazon basin lowlands, and the Galápagos Islands due to the different ecosystem that show each of these regions. The analysis was conducted in Arlequin 2.0 (Schneider et al. 2000). AMOVA test was also applied to find out if there were differences among *Ae. aegypti* populations from the three distinctive sampled regions (Galápagos, Pacific Coast and Amazon basin).

The haplotype sequences obtained were compared with the COI (971) and ND4 (509) sequences deposited in the GenBank from the region (Americas) and countries from Africa and Asia. The TCS network method helped determine the more closely related sequences (40 for COI and 45 for ND4) which were then used to infer the phylogenetic relationships with extra-Ecuadorian *Ae. aegypti* specimens with Bayesian inference analysis.

## Results

### Haplotypes distribution and frequency

The analysis of fragments of the genes COI (672 bp) and ND4 (262 bp) from 154 individuals of *Aedes aegypti* showed two haplotypes (H1, H2) present in Ecuador mainland and the Galápagos Islands. The phylogenetic analysis of the two detected mitochondrial haplotypes detected identified two well-supported clades. The COI-H1 was the most common in 60.4% of the specimens, while the H2 was detected in 39.6% of the specimens (Table 2). COI-H1 was detected in 14 localities, and H2 in nine localities, and both haplotypes were detected in seven localities, out of the 16 sampled localities (Table 2). The ND4-H1 was the most common, it was found in 62.3% of the specimens, while ND4-H2 was detected in 37.7% of the specimens (Table 3). ND4-H1 was detected in 17 localities, while ND4-H2 was detected in eleven localities, and both haplotypes (ND4-H1, ND4-H2) were detected in nine localities (Table 3). ND4 haplotypes showed eight polymorphic sites, all transitions at positions 31, 91, 100, 109, 148, 193, 226 and 241, with a length 262 bases (Fig. 2). The average nucleotide composition was 42.75% T, 31.68% A, 19.08% G, and 6.49% C, with a G+C content of 26.3%. COI haplotypes showed 13 polymorphic sites, all transitions (C - T or G - A) at positions 38, 173, 239, 242, 251, 323, 464, 473, 485, 584, 635, 641 and 665, with a length of 672 bases (Fig. 2). The average nucleotide composition was 39.14% T, 28.27% A, 17.11% C, and 15.48% G, with a G+C content of 32.9%. COI haplotype and nucleotide diversity of the entire data set (53 sequences) were Hd= 0.488 and π= 0.00943, respectively (Table 2). The average number of nucleotide differences k= 6.33962, and 13 segregating sites (S) were identified. Tajima’s test showed a significant value D= 3.60131 (p <0.05), and Fu and Li’s F*= 2.62213, D*= 1.51245 (Table 4). ND4 haplotype and nucleotide diversity of the entire data set (132 sequences) were Hd= 0.47 and π= 0.01447, respectively (Table 3). The average number of nucleotide differences k= 3.76313, and eight segregating sites (S) were identified. Tajima’s test showed a significant value D= 3.64936 (p <0.05), Fu and Li’s values F*= 2.4672 were significant, while D*= 1.23977 was not significant (Table 5). These values for COI and ND4 suggest balancing selection for both genes. The results indicate genetic variation without geographical restriction.

**Fig. 2.**
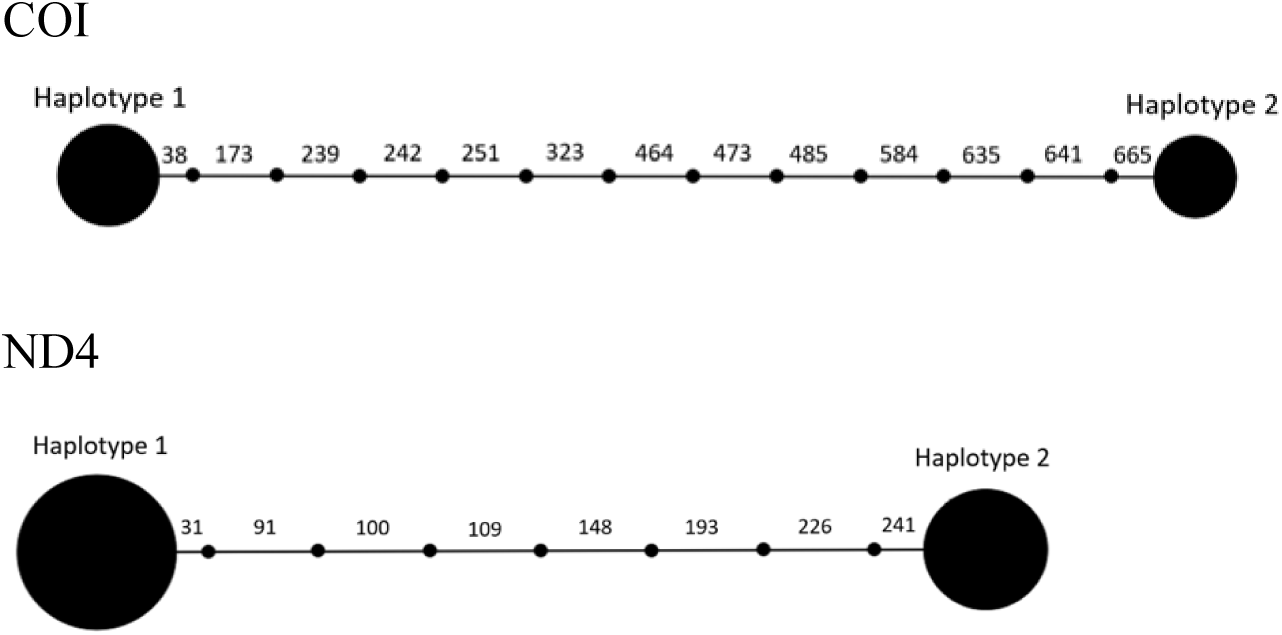
Haplotype network of COI and ND4 genes in *Aedes aegypti* populations from Ecuador. Numbers represent the mutational steps. The area of the circles is proportional to the frequency of each haplotype.

**Table 2.**
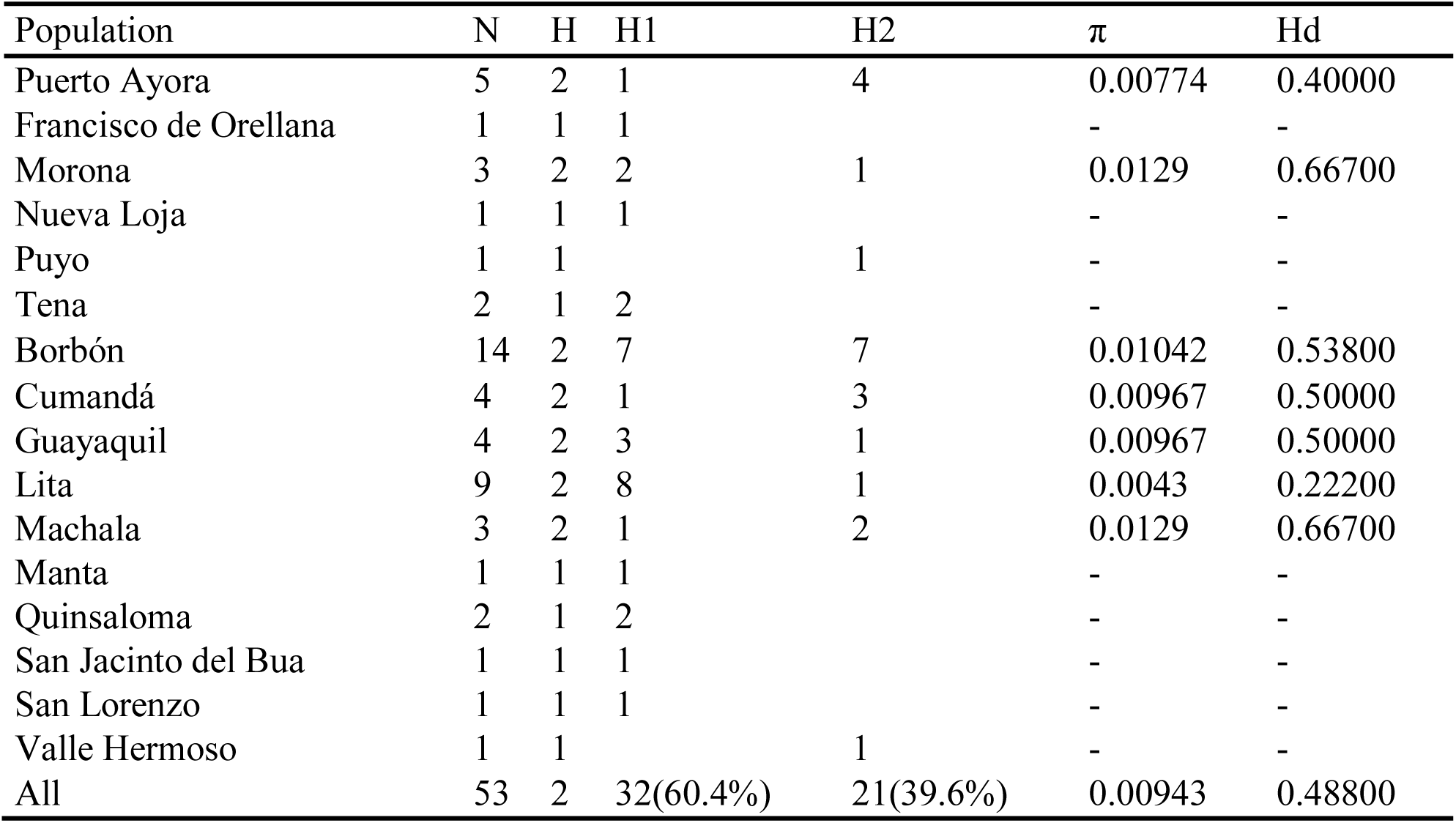
Polymorphism indexes of Aedes aegypti COI gene of 16 Ecuadorian populations. N: Sample size; H: haplotype per population; H1: haplotype 1 frequency: H2: haplotype 2 frequency π: nucleotide diversity; Hd: haplotype diversity

**Table 3.** Polymorphism indexes of ND4 gene of 19 Ecuadorian populations of *Aedes aegypti*. N: Sample size; H: haplotype per population; H1: haplotype 1 frequency: H2: haplotype 2 frequency π: nucleotide diversity; Hd: haplotype diversity.

| Population | N | H | H1 | H2 | $\pi$ | Hd |
| --- | --- | --- | --- | --- | --- | --- |
| Puerto Baquerizo Moreno | 3 | 1 | 3 |  | - | - |
| Puerto Ayora | 8 | 2 | 3 | 5 | 0.01636 | 0.53600 |
| Francisco de Orellana | 4 | 2 | 2 | 2 | 0.02036 | 0.66700 |
| Macas | 6 | 1 |  | 6 | - | - |
| Nueva Loja | 8 | 2 | 6 | 2 | 0.01309 | 0.42900 |
| Puyo | 3 | 1 | 3 |  | - | - |
| Tena | 5 | 1 | 5 |  | - | - |
| Babahoyo | 5 | 2 | 3 | 2 | 0.01832 | 0.60000 |
| Borbón | 10 | 2 | 6 | 4 | 0.01628 | 0.53300 |
| Cumandá | 8 | 2 | 3 | 5 | 0.01648 | 0.53600 |
| Esmeraldas | 5 | 2 | 2 | 3 | 0.01832 | 0.60000 |
| Guayaquil | 39 | 2 | 27 | 12 | 0.01335 | 0.43700 |
| Lita | 8 | 1 | 8 |  | - | - |
| Machala | 7 | 2 |  | 7 | - | - |
| Manta | 4 | 1 | 4 |  | - | - |
| Portoviejo | 1 | 1 | 1 |  | - | - |
| Quinsaloma | 2 | 1 | 2 |  | - | - |
| Santo Domingo | 3 | 2 | 2 | 1 | 0.02036 | 0.66700 |
| San Lorenzo | 1 | 1 | 1 |  | - | - |
| All | 130 | 2 | 81(62.3%) | 49(37.7%) | 0.01456 | 0.47300 |

**Table 4.** Neutrality test of Aedes aegypti COI gene of five Ecuadorian populations. A: P > 0.10 not significant; b: P<0.05 significant.

| Population |  | Tajima's<br>D | Fu & Li's<br>F* | Fu & Li's<br>D* |
| --- | --- | --- | --- | --- |
| Puerto Ayora | 5 | -1.21039<sup>a</sup> | -1.21039<sup>a</sup> | -1.21039<sup>a</sup> |
| Borbón | 14 | 2.86261<sup>b</sup> | 1.93118<sup>b</sup> | 1.4788<sup>b</sup> |
| Cumandá | 4 | -0.84307<sup>a</sup> | -0.85632<sup>a</sup> | -0.84307<sup>a</sup> |
| Guayaquil | 4 | -0.84307<sup>a</sup> | -0.84307<sup>a</sup> | -0.84307<sup>a</sup> |
| Lita | 9 | -1.88947<sup>b</sup> | -2.33051<sup>b</sup> | -2.14486<sup>b</sup> |
| All | 53 | 3.60131<sup>b</sup> | 2.62213<sup>b</sup> | 1.51245<sup>b</sup> |

**Table 5.** Neutrality test of ND4 gene of eight Ecuadorian populations of *Aedes aegypti*. a: P > 0.10 not significant; b: P<0.05 significant.

| Population |  | Tajima's<br>D | Fu & Li's<br>F* | Fu & Li's<br>D* |
| --- | --- | --- | --- | --- |
| Puerto Ayora | 8 | 1.88129<sup>a</sup> | 1.7279<sup>b</sup> | 1.46524<sup>b</sup> |
| Francisco de Orellana | 4 | 2.19753<sup>a</sup> | 2.16298<sup>a</sup> | 2.19753<sup>b</sup> |
| Nueva Loja | 8 | 0.53786<sup>a</sup> | 1.38232<sup>a</sup> | 1.46524<sup>b</sup> |
| Babahoyo | 5 | 1.76148<sup>a</sup> | 1.84469<sup>a</sup> | 1.76148<sup>b</sup> |
| Borbón | 10 | 2.19508<sup>b</sup> | 1.80255<sup>b</sup> | 1.41121<sup>b</sup> |
| Cumandá | 8 | 1.88129<sup>a</sup> | 1.7279<sup>b</sup> | 1.46524<sup>b</sup> |
| Esmeraldas | 5 | 1.76148<sup>a</sup> | 1.84469<sup>a</sup> | 1.76148<sup>b</sup> |
| Guayaquil | 39 | 2.43323<sup>b</sup> | 1.95088<sup>b</sup> | 1.31228<sup>a</sup> |
| All | 130 | 3.67790<sup>b</sup> | 2.4672<sup>b</sup> | 1.23977<sup>a</sup> |

COI haplotype diversity of the individual populations varied from 0.222 (Lita) to 0.667 (Machala and Morona). Nucleotide diversity by locality (π) varied from 0.0043 (Lita) to 0.0219 (Morona and Machala) (Table 2). Neutrality tests (D, F*, D*) for all samples (53) were positive and significant (Table 4). Neutrality tests (D, F*, D*) showed positive values and were significant for sequences from Borbón, while Lita sequences analysis showed negative values and were significant. The rest of localities showed negative values that were not significant (Puerto Ayora, Cumandá and Guayaquil) (Table 4).

ND4 haplotype diversity (Hd) of the populations varied from 0.429 (Nueva Loja) to 0.667 (Francisco de Orellana, Sto. Domingo) (Table 3). While the nucleotide diversity (π) in the sampled locations varied from 0.01309 to 0.02036. Fu & Li’s D* values were positive and significant for all locations, except Guayaquil (Table 5). Tajima’s D and Fu & Li’s F* for all the sequences were positive and significant, while Fu & Li’s D* was positive and non-significant. Tajima neutrality test (D) showed positive values that were significant (Borbón, Guayaquil), the rest of values were positive and not significant. Fu and Li’s F* values were positive and significant for some locations (Puerto Ayora, Cumandá, Borbón and Guayaquil). The D* values were significant for all locations, except Guayaquil (Table 5).

AMOVA test applied to the whole data set of the COI individual gene sequences showed COI Fixation Index (F_ST_= 0.12328), which was not significant (Table 6). The variation within populations was greater (87.7%) than the variation among populations (12.3%) (Table 6). On the other hand, AMOVA test applied to the whole data set of the ND4 gene sequences showed a Fixation Index F_ST_ = 0.22668, which was significant. The variation within populations was greater (77.3%) that the variation among populations (22.7%) (Table 7).

**Table 6.**
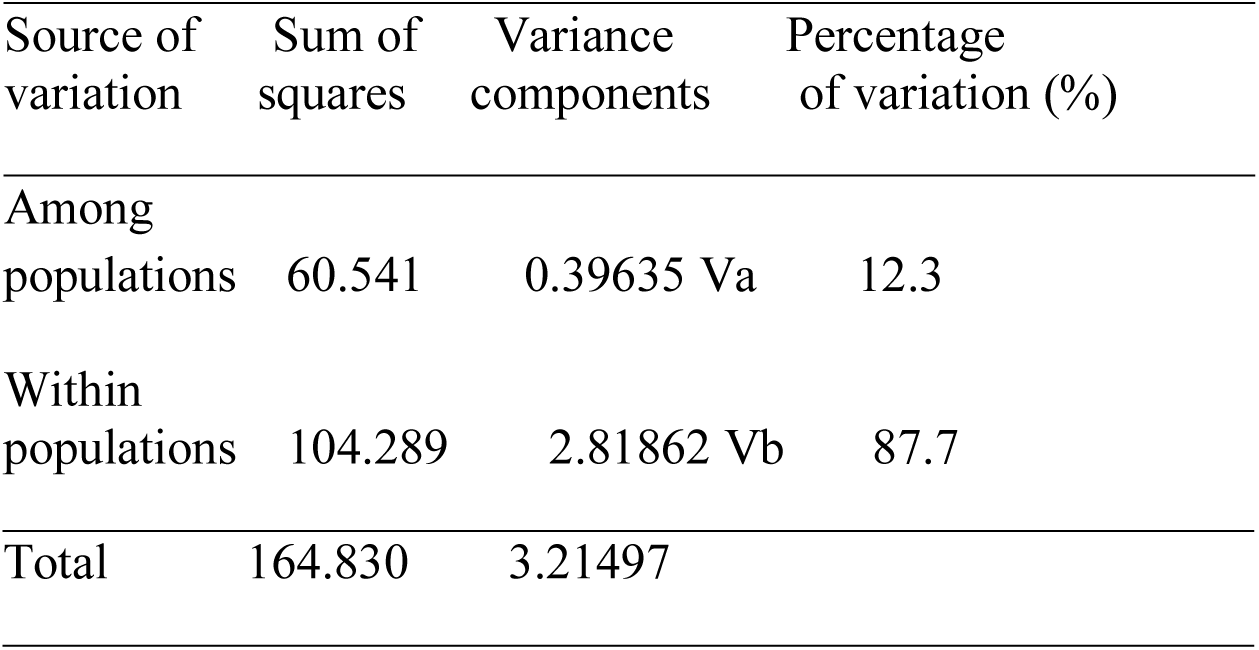
AMOVA analysis of COI sequences of Aedes aegypti from five locations in Ecuador. Fixation Index FST= 0.12328, p-value = 0.13294+-0.01173. Not significant, p-value > 0.10

**Table 7.** AMOVA analysis of ND4 haplotypes of *Aedes aegypti* from five locations in Ecuador. Fixation Index FST= 0.22688. P value < p (0.05) significant (Va, Vb).

| Source of variation | Sum of squares | Variance components | Percentage of variation |
| --- | --- | --- | --- |
| Among populations | 77.690 | 0.43835 Va | 22.7 |
| Within populations | 168.795 | 1.49376 Vb | 77.3 |
| Total | 246.485 | 1.93211 |  |

The analysis comparing the sequences by geographical region showed COI haplotype diversity of Galápagos was 0.4, Pacific Coast 0.481 and Amazon basin 0.429. Nucleotide diversity varied 0.00774 (Galápagos), 0.093 (Pacific coast) and 0.00829 (Amazon basin) (Table 8). Neutrality tests (Tajima D, Fu & Li F*) showed negative values for Galápagos and D* (Fu & Li) was positive, all the vales were not significant (Table 9). Neutrality tests (D, F*, D*) for sequences from the Pacific coast were positive and significant. While for the Amazon basin values (D, F*, D*) were positive and only Tajima’s D* was significant. AMOVA test among *Ae. aegypti* populations separating the samples from the three distinctive sampled regions (Galápagos, Pacific Coast and Amazon basin) showed not significant Fixation indices (F_ST_, F_SC_ and F_CT_) (Table 10).

**Table 8.** Polymorphism indexes of of Aedes aegypti COI gene from three geographical regions in Ecuador. N: Sample size; H: haplotype per population; H1: haplotype 1 frequency: H2: haplotype 2 frequency π: nucleotide diversity; Hd: haplotype diversity

| Region | N | H | H1 | H2 | $\pi$ | Hd |
| --- | --- | --- | --- | --- | --- | --- |
| Galápagos | 5 | 2 | 1 | 4 | 0.00774 | 0.40000 |
| Amazon basin | 8 | 2 | 6 | 2 | 0.00829 | 0.42900 |
| Pacific coast | 40 | 2 | 25 | 15 | 0.00930 | 0.48100 |

**Table 9.**
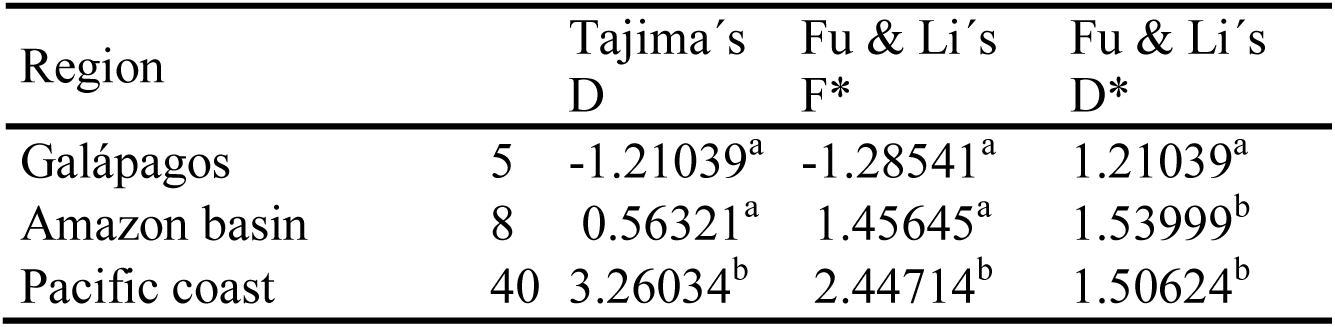
Neutrality test of *Aedes aegypti* COI gene from three geographical regions in Ecuador. a: P > 0.10 not significant; b: P<0.05 significant.

**Table 10.** AMOVA analysis of COI gene sequences of Aedes aegypti from three geographical regions in Ecuador. Fixation Indices FST= 0.15727, FSC= 0.09227, FCT= 0.07161. P-values were not significant (Vb, Vc). Significance tests (1023 permutations).

| Source of variation | Sum of squares | Variance components | Percentage of variation |
| --- | --- | --- | --- |
| Among groups | 13.055 | 0.23951 Va | 7.1 |
| Among populations within groups | 47.486 | 0.28650 Vb | 8.5 |
| Within populations | 104.289 | 2.81862 Vc | 84.2 |
| Total | 164.830 | 3.34463 |  |

ND4 Gene haplotype diversity in Galápagos sample was 0.545, Pacific Coast 0.469 and Amazon basin 0.484. Nucleotide diversity for sequences from Galápagos was 0.01666, 0.01443 for the Pacific coast and 0.01490 for the Amazon basin sequences (Table 11). Neutrality tests (D, F*) were positive values and significant, while D* value was positive and significant only for Galápagos (Table 12). Fixation Index (F_ST_) value was 0.18728 that was significant, F_SC_ was 0.26360 and significant, while F_CT_ was negative (−010365) and was not significant (Table 13). Negative values indicate excess of heterozygotes and should be interpreted as zero in the AMOVA (Schneider et al. 2000).

**Table 11.** Polymorphism indexes of ND4 gene of *Aedes aegypti* from three geographical regions in Ecuador. N: Sample size; H: haplotype per population; H1: haplotype 1 frequency: H2: haplotype 2 frequency π: nucleotide diversity; Hd: haplotype diversity.

| Region | N | H | H1 | H2 | $\pi$ | Hd |
| --- | --- | --- | --- | --- | --- | --- |
| Galápagos | 11 | 2 | 6 | 5 | 0.01666 | 0.54500 |
| Amazon basin | 26 | 2 | 16 | 10 | 0.01490 | 0.48400 |
| Pacific coast | 93 | 2 | 59 | 34 | 0.014430 | 0.46900 |

**Table 12.**
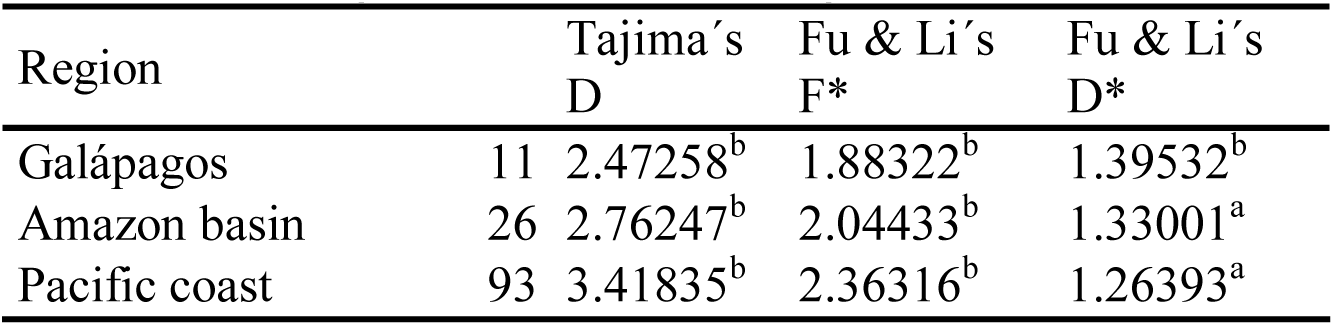
Neutrality test of ND4 gene of Aedes aegypti from three geographical regions in Ecuador. a: P > 0.10 not significant; b: P<0.05 significant.

**Table 13.** AMOVA analysis of ND4 gene sequences of *Aedes aegypti* from three geographical regions in Ecuador. Fixation Indices F_ST_= 0.18728, F_SC_= 0.26360, F_CT_= − 0.10365. P-values were not significant (Va, Vb, Vc). Significance tests (1023 permutations).

| Source of variation | Sum of squares | Variance components | Percentage of variation |
| --- | --- | --- | --- |
| Among groups | 0.679 | -0.19050 Va | -10.3 |
| Among populations within groups | 77.011 | 0.53471 Vb | 29.0 |
| Within populations | 168.795 | 1.49376 Vc | 81.2 |
| Total 131 | 246.485 | 1.83797 |  |

Combined analysis of both COI and ND4 genes of 29 sequences from ten localities resulted also in two haplotypes (Table 14). There were 21 sequences of haplotype 1 (H1) (73.3%) and 8 sequences of haplotype 2 (H2) (26.7%) (Table 14). H1 was detected in nine locations, H2 in five and both haplotypes (H1, H2) were present in four locations (Table 14). Haplotype diversity from all data set was 0.405 and nucleotide diversity was 0.0091. Haplotype diversity in analyzed locations varied from 0.5 (Borbón), to 1.0 (Santa Cruz, Cumandá and Guayaquil). Neutrality tests (D, F*, D*) for the whole data set were positive and significant (Table 15). AMOVA test applied to the whole data set (29 sequences) of the COI-ND4 genes sequences showed F_ST_= 0.12531 (Fixation Index), which was not significant. The variation within populations was greater (87.4%) that the variation among populations (12.5%) (Table 16).

**Table 14.** Polymorphism indexes of COI and ND4 concatenated gene sequences of ten Ecuadorian populations of Aedes aegypti. N: Sample size; H: haplotype per population; H1: haplotype 1 frequency: H2: haplotype 2 frequency π: nucleotide diversity; Hd: haplotype diversity.

| Population | N | H | Haplotype | H1 | H2 | $\pi$ | Hd |
| --- | --- | --- | --- | --- | --- | --- | --- |
| Puerto Ayora | 2 | 2 | I, II | 1 | 1 | 0.02248 | 1 |
| Tena | 2 | 1 | I | 2 |  | - | - |
| Borbón | 9 | 1 | I, II | 6 | 3 | 0.01124 | 0.5 |
| Cumandá | 2 | 2 | I, II | 1 | 1 | 0.02248 | 1 |
| Guayaquil | 2 | 2 | I, II | 1 | 1 | 0.02248 | 1 |
| Lita | 6 | 2 | I | 6 |  | - | - |
| Machala | 2 | 2 | I, II |  | 2 | - | - |
| Manta | 1 | 2 | I | 1 |  | - | - |
| Quinsaloma | 2 | 2 | I | 2 |  | - | - |
| San Lorenzo | 1 | 2 | I | 1 |  | - | - |
| All | 29 | 2 | I, II | 21(73.3%) | 8 (26.7%) | 0.0091 | 0.405 |

**Table 15.**
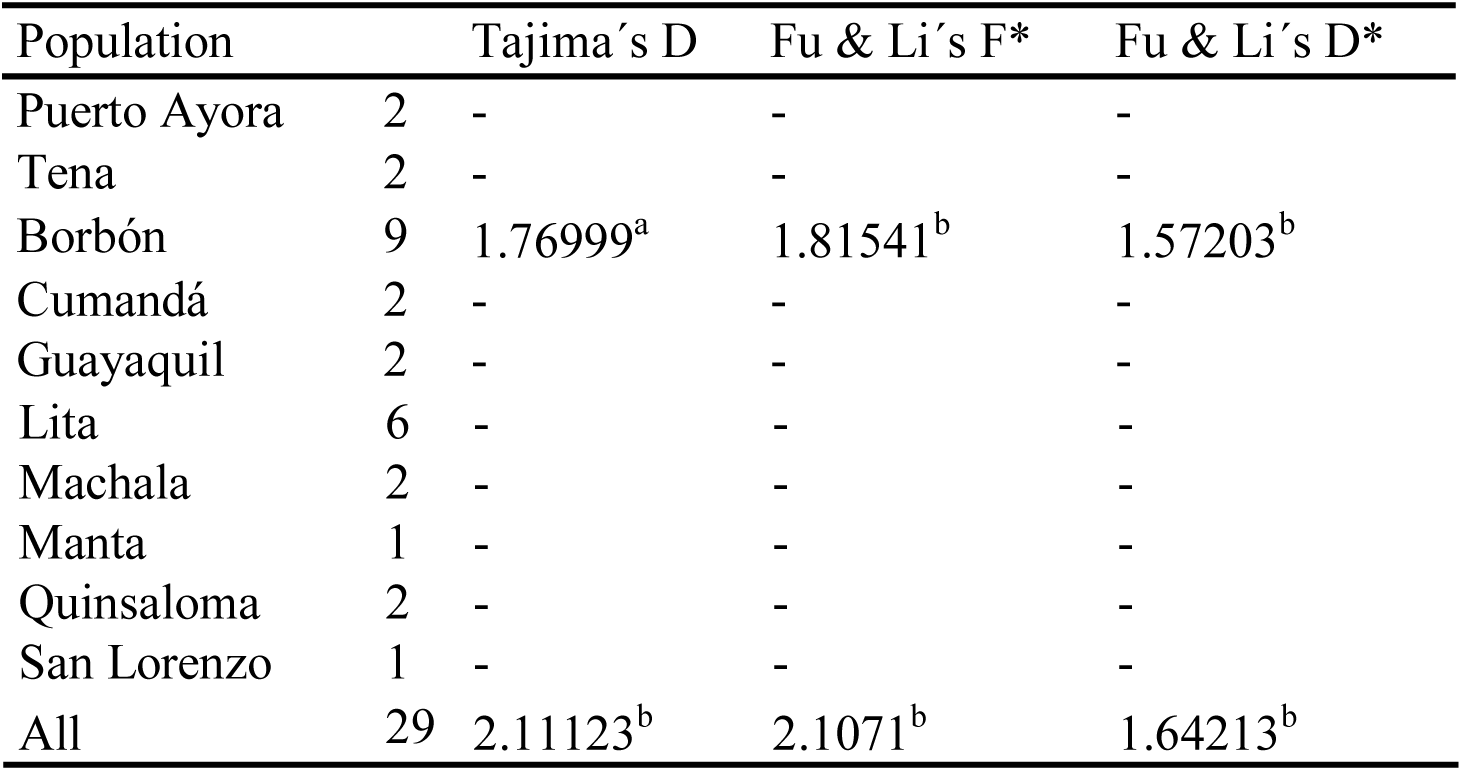
Neutrality test of COI and ND4 concatenated genes of *Aedes aegypti*. a: P > 0.10 not significant; b: P<0.05 significant.

**Table 16.** AMOVA analysis of concatenated sequences of COI and ND4 concatenated gene sequences of Aedes aegypti in ten locations in Ecuador. Fixation Index F_ST_= 0.12531. P-value was not significant. Significance tests (1023 permutations).

| Source of variation | d.f. | Sum of squares | Variance components | Percentage of variation |
| --- | --- | --- | --- | --- |
| Among populations | 9 | 48.155 | 0.55422 Va | 12.5 |
| Within populations | 19 | 73.500 | 3.86842 Vb | 87.4 |
| Total | 28 | 121.655 | 4.42264 |  |

COI haplotype network showed 13 mutational steps, while the ND4 haplotype network showed eight mutational steps (Fig. 2). The size of the circle represents the occurrence of each haplotype. The low frequency of H2 might suggest that is a derivation from H1.

Phylogenetic analysis of COI and ND4 sequences showed two haplotypes. The grouping of the locations did not show a geographic pattern (Figs. 3, 4). Combined phylogenetic analysis of concatenated COI and ND4 sequences showed also two haplotypes, the same as the individual analysis of both genes. COI-ND4-H1 was the most abundant in the 10 analyzed localities (Tables 2, 3). There was also coincidence in the grouping of all COI and ND4 sequences when analyzed individually and also when both genes were concatenated and analyzed (Fig. 5). The analysis of concatenated COI and ND4 (29 sequences) by geographical region (Galápagos, Amazon basin and Pacific coast) resulted in haplotype diversity (Hd) of 1.0 for Galápagos, and 0.420 for the Pacific coast, nucleotide diversity (π) 0.02248 for Galápagos and 0.00944 for the Pacific coast (Table 17). Tajima’s test (D statistic) and Fu and Li’s (F* and D*) showed positive significant values (D= 2.12789 (p <0.05), F*= 2.07874 and D*= 1.61887). (Table 18). The Fixation Indices (F_ST_= −0.00901, F_SC_= 0.17210, F_CT_= 0.21875) were not significant. The percentage of variation within populations was the highest (100.9%) (Table 19).

**Fig. 3.**
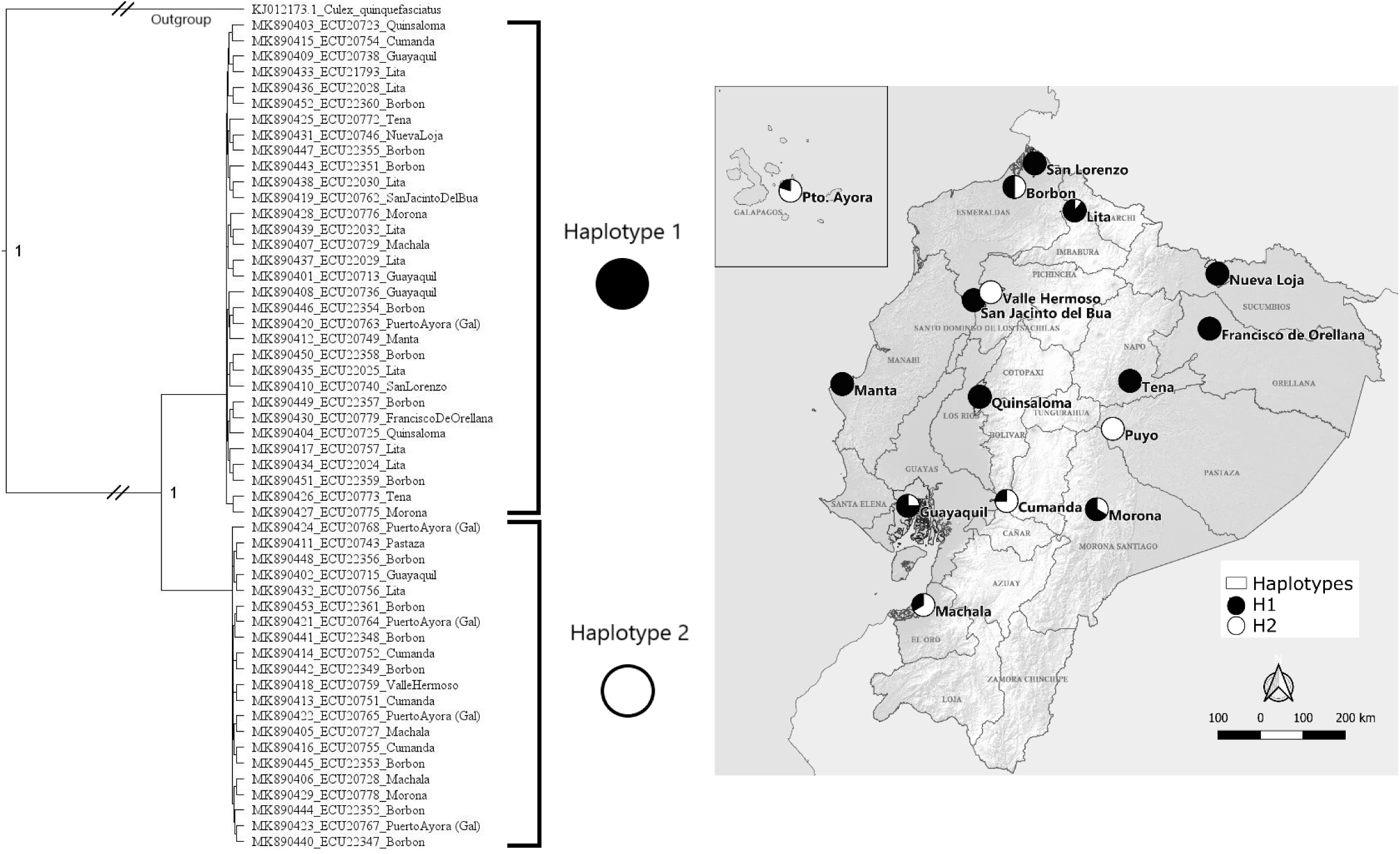
Phylogenetic analysis of COI gene sequences of *Aedes aegypti* from Ecuador

**Fig. 4.**
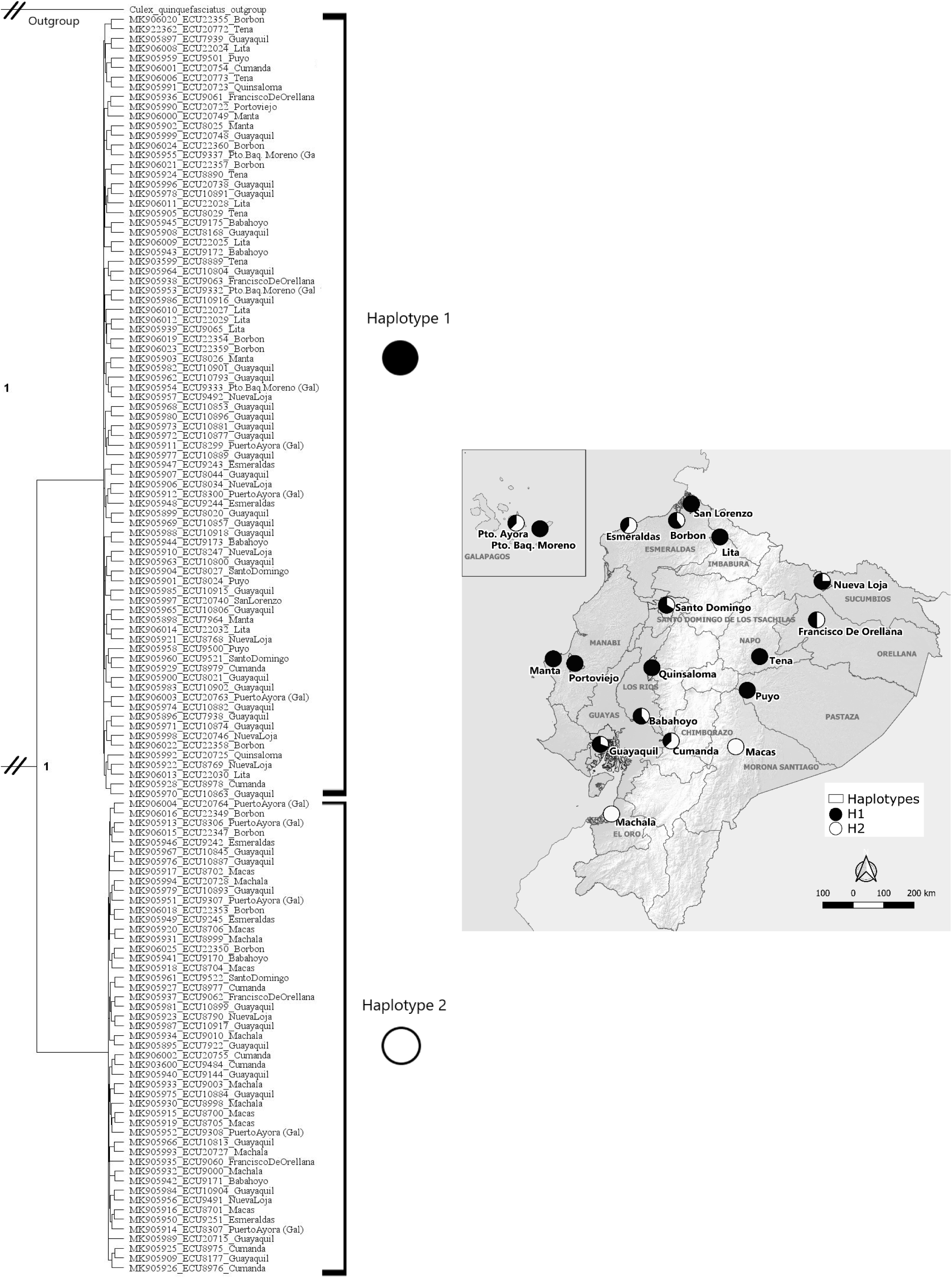
Phylogenetic analysis of ND4 gene sequences of *Aedes aegypti* from Ecuador

**Fig 5.**
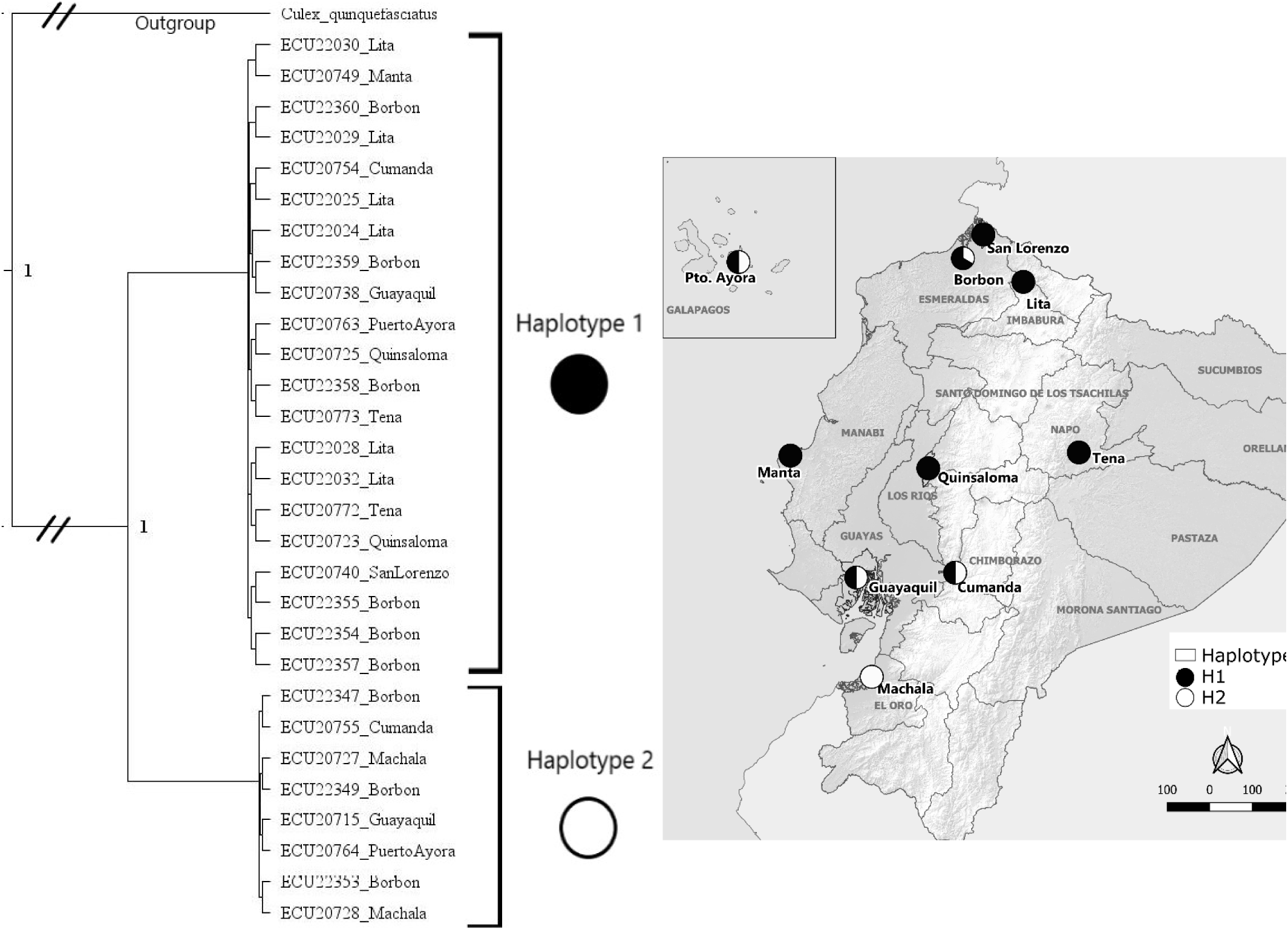
Phylogenetic analysis of COI and ND4 concatenated gene sequences of *Aedes aegypti* from Ecuador

**Table 17.** Polymorphism indexes of COI and ND4 concatenated genes of *Aedes aegypti* from three geographical regions in Ecuador. N: Sample size; H: haplotype per population; H1: haplotype 1 frequency: H2: haplotype 2 frequency π; nucleotide diversity; Hd: haplotype diversity

| Region | N | H | H1 | H2 | $\pi$ | Hd |
| --- | --- | --- | --- | --- | --- | --- |
| Galápagos | 2 | 2 | 1 | 1 | 0.02248 | 1.000 |
| Amazon basin | 2 | 1 | 2 |  | - | - |
| Pacific coast | 25 | 2 | 18 | 7 | 0.00944 | 0.420 |

**Table 18.**
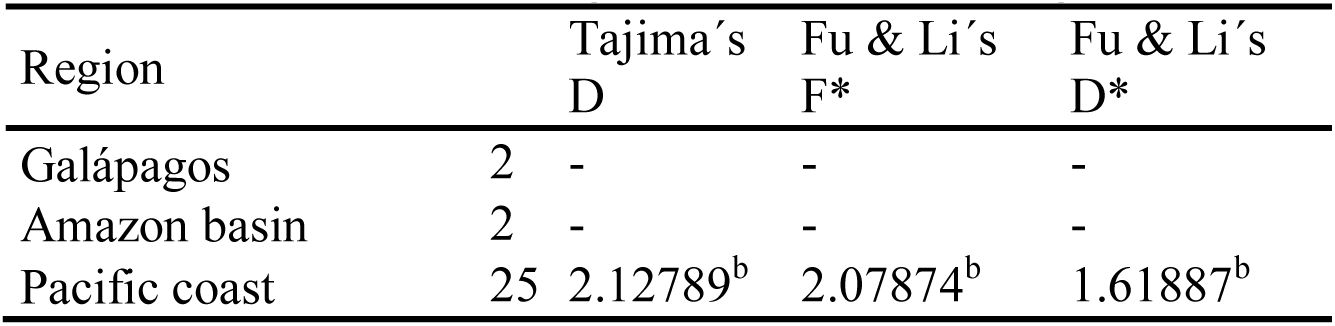
Neutrality test of COI and ND4 concatenated genes of Aedes aegypti from three geographical regions in Ecuador. a: P > 0.10 not significant; b: P<0.05 significant.

**Table 19.** AMOVA analysis of COI and ND4 concatenated gene sequences of *Aedes aegypti* from three geographical regions in Ecuador. Fixation Indices FST= −0.00901, FSC= 0.17210, FCT= 0.21875. P-values were not significant (Va, Vb, Vc). Significance tests (1023 permutations).

| Source of variation | Sum of squares | Variance components | Percentage of variation |
| --- | --- | --- | --- |
| Among groups | 5.315 | -0.83867 Va | -21.8 |
| Among populations within groups | 42.840 | 0.80414 Vb | 20.9 |
| Within populations | 73.500 | 3.86842 Vc | 100.9 |
| Total | 121.655 | 3.83388 |  |

COI-H1 (ECU22032_Lita) grouped with sequences from Colombia, Brazil, Bolivia (Americas); Benin and Guinea (West and Central Africa), Kenya (East Africa), India (Asia) and Australia. While COI-H2 (ECU20764_Pto. Ayora) grouped with sequences from Sri Lanka, India, Cambodia, Thailand (Asia); Mexico, Brazil, USA (Americas) and close to sequences from Martinique and USA (0.9207) (Fig. 6). ND4-H1 (ECU22025_Lita) grouped with similar sequences from USA, Mexico, Colombia, Bolivia, Brazil, Perú, Chile (Americas); Myanmar (Asia); Ivory Coast, Guinea, Nigeria, Cameroon, Senegal (West Africa). ND4-H2(ECU8306_Pto.Ayora) grouped with sequences from Mexico, Brazil, Colombia and Peru (Americas) (Fig. 7).

**Fig. 6.**
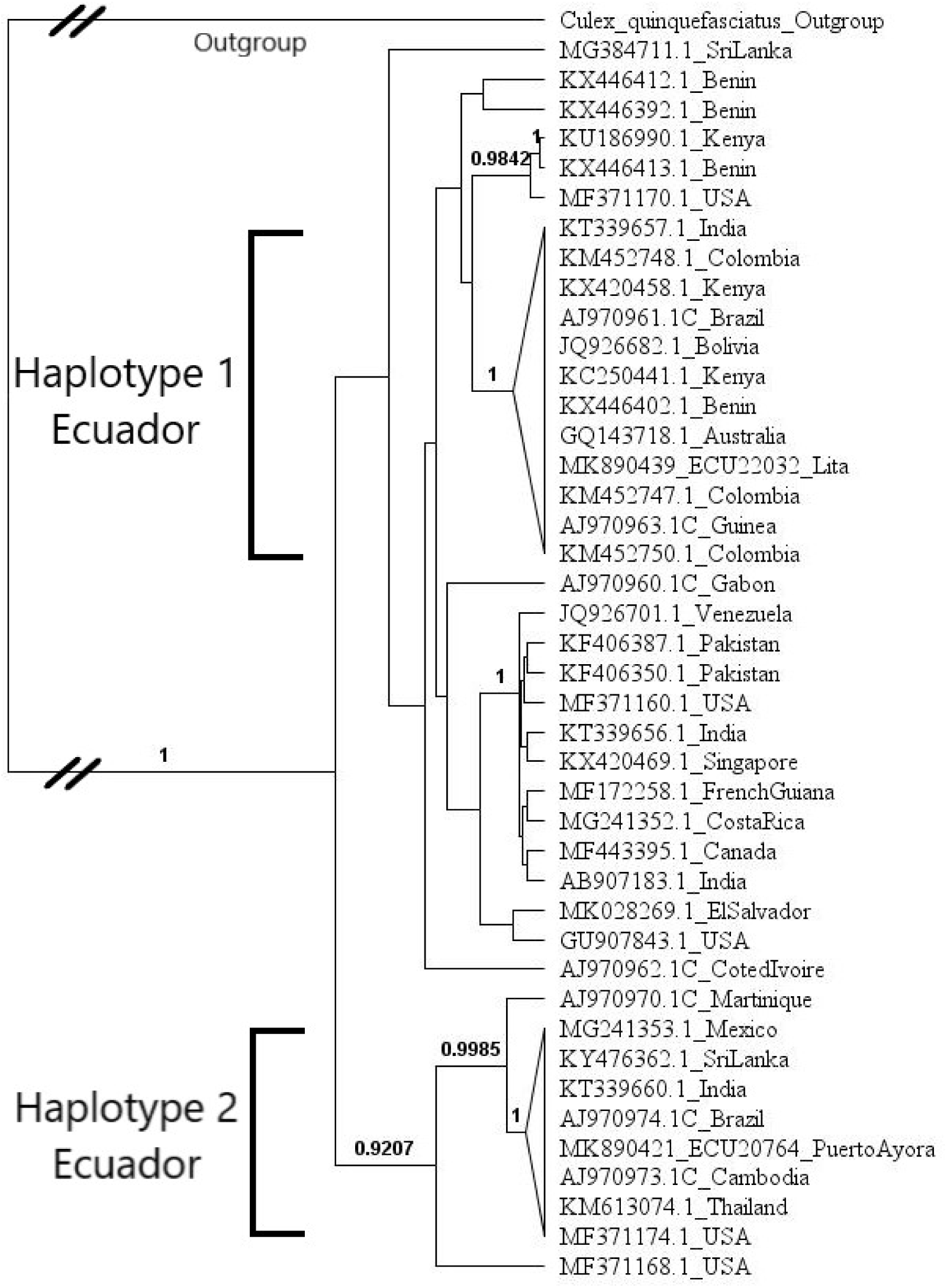
Phylogenetic tree showing the grouping of haplotypes (COI gene) of *Aedes aegypti* from Ecuador with sequences from Africa, Asia and the Americas.

**Fig. 7.**
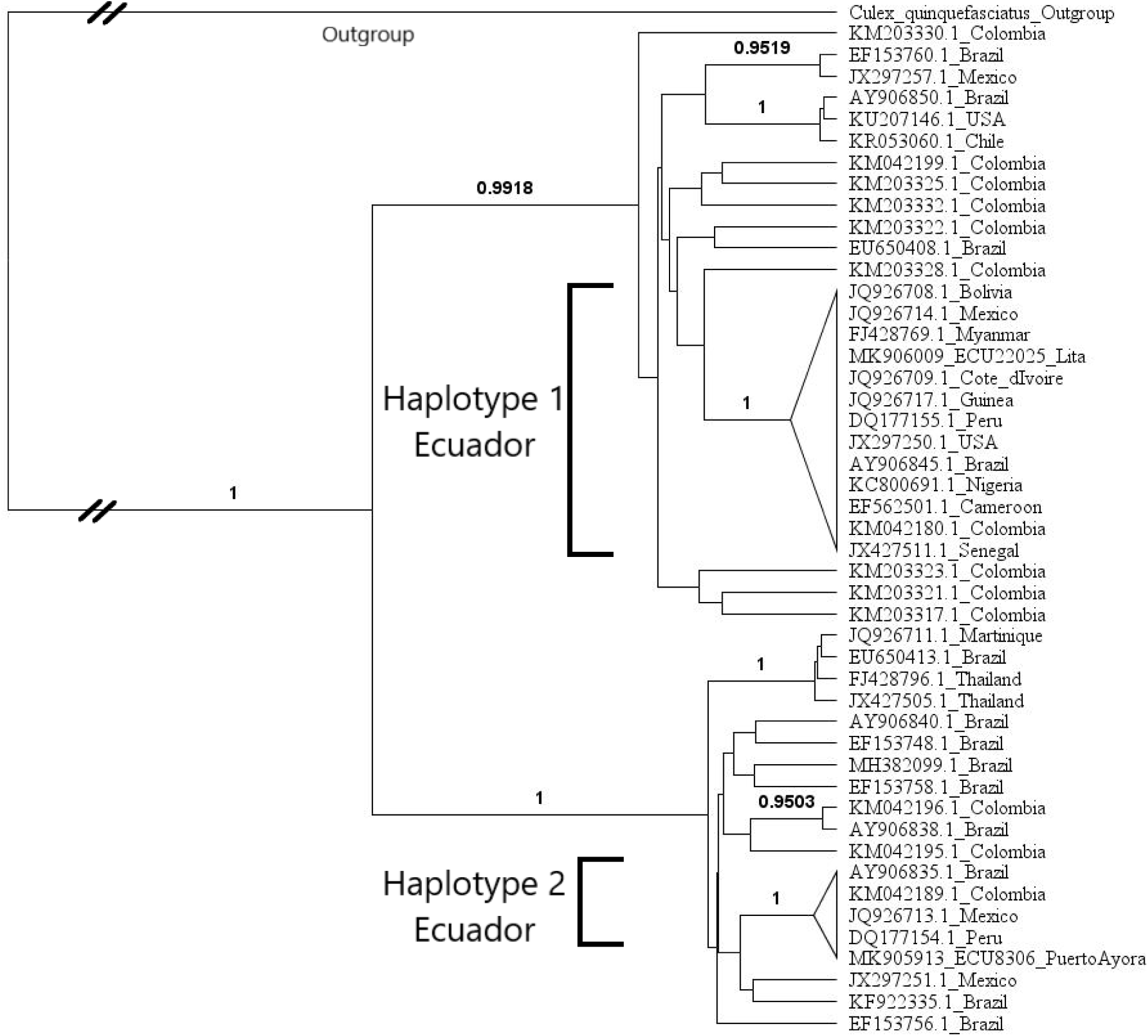
Phylogenetic tree showing the grouping of haplotypes (ND4 gene) of *Aedes aegypti* from Ecuador with sequences from Africa, Asia and the Americas.

We report overlapping peaks in four sequences (Machala, Lita, Macas and Francisco de Orellana) that indicate the presence of two nucleotides at a single site, this suggests heteroplasmy in the individuals.

## Discussion

This is the first report on the genetic variation of *Aedes aegypti* populations in Ecuador. *Aedes aegypti* apparently arrived in Ecuador during the late 18^th^ century, by the beginning of the 19^th^ century Guayaquil city in the Pacific coast was considered a main focus of yellow fever presumably transmitted by this mosquito (Kassirsky and Plotnikov 2003). Since 1988 the mosquito has been responsible of important arboviral outbreaks with the presence of four serotypes of Dengue virus (DENV), Zika (ZIKV) Chikungunya (CHIKV) and presumably also the Yellow fever virus (YFV) (Brathwaite-Dick et al. 2012). However, since the re-infestation of the vector in the late 1980s the mosquitoes have extended from the Ecuadorian Pacific coastal region to the Amazon basin region, reaching locations at 1,650 m of altitude at the northeastern mountain slopes.

Several studies have shown the genetic variation of *Ae. aegypti* populations in the neighboring countries, Perú (Costa-da-Silva et al. 2005), Colombia (Jaimes-Dueñez et al. 2015, Atencia et al. 2018) and Brazil (Monteiro et al. 2014, Paduan and Ribolla 2008). In Ecuador, there were two haplotypes, which show low genetic diversity in all the sampled localities, although the ecological differences among the geographical areas are apparent among the Pacific coast, Amazon basin and the Galápagos Islands. Both (H1, H2) haplotypes are present in all the geographical areas, although the H2 is less common. This may suggest that regardless of the genetic structure of the mosquito populations, they are able to disperse and get established in all suitable areas.

The phylogenic trees indicate that *Ae. aegypti* COI-H1 from Ecuador are grouped with mosquitoes from the Americas, West, Central and East Africa, Asia and Australia. In the case of COI-H2 includes sequences from Asia and the Americas. The ND4-H1sequences are similar to sequences from the Americas, Asia and West Africa. While the ND4-H2 sequences grouped with sequences from the Americas. Overall, the H1 group of *Ae. aegypti* seems to be related to populations from the West African while H2 is related to the Asian populations. This suggest that *Ae. aegypti* arrived in Ecuador originally from Africa to Asia and afterwards to the Americas, the same pattern reported for ancestral populations (Bracco et al. 2007, Costa-da-Silva et al. 2005, Paupy et al. 2012). *Aedes aegypti* may have arrived in Ecuador through the seaports in the Pacific coast, especially Guayaquil, as it suggests the first detection of *Aedes albopictus* in this city (Ponce et al. 2018). *Aedes aegypti* apparently spreads from the Pacific lowlands to Galápagos and to the Amazon basin lowlands, and this process may be constant and dynamic due to human mobility and goods trade that promote the dispersal of the mosquito. Although Galápagos applies strict touristic and resident regulations*, Ae. aegypti* has been reported in the main inhabited islands since 2001 (Causton et al 2006). The introduction of *Ae. aegypti* has caused dengue outbreaks in Puerto Ayora (Santa Cruz) and Puerto Baquerizo Moreno (San Cristóbal) cities (Real-Cotto et al. 2017).

Our analysis showed low nucleotide diversity for COI (0.00943) and ND4 gene (0.01456), which indicates that genetic diversity of *Ae. aegypti* populations is low. The low level of genetic diversity observed in Ecuadorian cities may be the result of mosquito control activities combined with reduced number of mosquitoes during the dry season. These factors may induce intraspecific inbreeding and as result low genetic diversity. Previous studies reported in Brazil (Bracco et al. 2007), Venezuela (Herrera et al. 2006), and Mexico (Gorrochotegui et al. 2002) showed high nucleotide diversity (π = 0.0199, 0.01877 and 0.02161 respectively). These authors also suggest a significant correlation between gene flow in *Ae. aegypti* and human transportation, explaining the similar genetic composition between Ecuadorian and American populations of this species. The similarity among the American populations could be the result of constant gene flow and reinvasion processes (Bracco et al. 2007). This hypothesis supports the idea of demographic expansion and high level of colonization in all the Ecuadorian regions including Galápagos Islands. However, it is remarkable that in the neighboring countries there are reports of several haplotypes. The variability observed may also be affected by the use of insecticide control. Although the use of insecticides is expected to produce high levels of genetic differentiation as reported in Brazil (Ayres et al. 2004). It may also be a selection pressure that may be favoring the spread of only the two haplotypes detected. In Ecuador, the mosquito has been subjected to selection pressure by intensive and extensive adult (malathion and deltamethrin) and immature (temephos) insecticide use (Morales et al. 2019), that may cause bottleneck selection and consequently low genetic variability. Random amplified polymorphic DNA (RAPD) analysis of Brazilian *Ae. aegypti* populations showed high heterozygosity in areas treated with insecticides, apparently this condition is the result of a combination of variation of population densities, spatial heterogeneity, and intense insecticide treatment (Ayres et al. 2004). The factors that maintain the low diversity population structure of *Ae. aegypti* in our sampled areas need to be determined, since differences in the genetic structure are in relation to ecological conditions observed in French Polynesia (Paupy et al. 2000), Thailand (Mousson et al. 2002), Vietnam (Huber et al. 2002), Brazil (Ayres et al. 2003, Ayres et al. 2004). Nucleotide diversity value (π) in the ND4 mitochondrial gene was higher than in the mitochondrial COI gene. According to Gorrochotegui et al. (2002) the difference may be due to higher constraints on the mutation rate in the COI gene.

The phylogenetic results show similarity with mitochondrial haplotypes of COI gene previously reported in Latin America (Fraga et al. 2013, Bracco et al. 2007, Costa-da-Silva et al. 2005, Paupy et al. 2012).

The closeness and human activities among Ecuador, Colombia and Peru, may easily explain the similarities of the shared lineages of ND4 gene, which are common among these countries. However, similar lineages have also been reported from several other countries in the Americas. On the other hand, COI sequences show that haplotype 1 share similarity with sequences from Asia and the Americas. While haplotype 2, apparently the most widely spread lineage reported, shares similarity with sequences from Asia, West and East Africa and the Americas. According to Eltis and Richardson (2010) *Ae. aegypti* populations reached the Americas about 440-550 years ago through slave trading from West Africa, therefore additional genetic data and analysis may be required to trace the spread of the *Aedes aegypti* lineages around the world. Powell and Tabachnick (2013) mention that *Ae. aegypti*, according to genetic evidence, probably came from West Africa into the New World, and then spread to Asia and Australia. Consequently, populations in the New World would be derived directly from African populations, while Asia/Australian populations are derived from New World populations. The genetic identity of *Ae. aegypti* may be a factor that affects the infection, susceptibility and capacity to transmit Dengue virus (DENV) and it is known that the mosquito genotype may modulate the transcriptional response depending on the strain of the DENV (Behura et al. 2014). The most common haplotype in high risk arbovirus areas in Ecuador is the H1, which may have relationship with a higher risk for virus transmission as described in Colombia (Jaimes-Dueñez et al. 2015). The haplotype 1 (H1) is the most abundant and may be related with the incidence of arboviral diseases in Ecuador. Souza-Neto et al. (2019) mentions that *Ae. aegypti* shows complete susceptibility to get infected by Zika (ZIKV), dengue (DENV) and chikungunya (CHIKV), but not yellow fever viruses (YFV). The number of dengue cases in Ecuador since year 2000 throughout 2015 shows that DENV serotypes has shifted from DENV-2 (year 2000) to DENV-3 (years 2001-2006), and DENV-1 and DENV-2 (years 2007-2015), with less cases of DEN4 (Real-Cotto et al. 2017). This epidemiological trend may suggest that Ecuadorian mosquito populations are more susceptible in descendant order to DENV-3, DENV-1, DENV-2 and DENV-4. In Brazil *Ae. aegypti* is particularly susceptible to DENV-2 (Souza-Neto et al. 2019). The apparent mixed presence of only two haplotypes in all geographic areas may have epidemiological, vector control and pest management implications. Calvez et al. (2016) demonstrated that infection and replication of ZIKV in *Ae. aegypti* also may differ due to vector and virus lineages (Sim et al. 2014). The Zika virus (ZIKV) circulating in Ecuador corresponds to the Asian lineage (Cevallos et al. 2018), which apparently is less infective to *Ae. aegypti* populations from the American continent than the African ZIKV lineage (Souza-Neto et al. 2019).

Monteiro et al. (2014) suggests that the two major genetic groups found in Brazil populations of *Ae. aegypti* are recolonizations after the eradication programs in the 60’s last century. Considering that the sequences detected in Ecuador are similar with some detected in Brazil, it may also be possible that *Ae. aegypti* populations in Ecuador were also recolonizations. However, grouping of the haplotypes detected in Ecuador with sequences from Asia and Africa and their distribution do not have a geographic pattern like the genetic groups in Brazil (Monteiro et al. 2014), that may indicate recolonization process. The absence of a defined spatial pattern of two genetic groups was also found in Colombian and Peruvian cities (Costa-da-Silva et al. 2005, Jaimes-Dueñez et al. 2015). Jaimes-Dueñez et al. (2015) also reports one haplotype widespread distributed like occurs with the H1 in Ecuador.

We also report the presence of four sequences for ND4 which contain both variations (H1 and H2) in the eight reported segregating sites, this corresponds to heteroplasmy. This condition occurs when there is more than one type of mitochondrial DNA in a cell or mitochondrion (Melton 2004). The presence of mitochondrial pseudogenes has been observed in nuclear and mitochondrial genome of *Ae. aegypti*, (Hlaing et al. 2009, Paduan and Ribolla 2008).

Heteroplasmic conditions have been reported in mammals, fish and insects (Wilkinson and Chapman 1991, Volz-Lingenhohl et al. 1992, Nesbø et al. 1998). In Drosophila, heteroplasmy has been suggested to be produced by a single mutation event and paternal leakage of mtDNA (Solignac et al. 1986, Satta and Chigusa 1991). It has been mentioned that this condition may be a contamination by excessive number of amplification cycles. However, all our samples were amplified following the same protocol. Heteroplasmy in humans is linked to maternal inherited diseases, however its effects in insects are unknown (Cataldo et al. 2013).

The GC content in both genes was low (COI= 26.3%, ND4= 32.9%), apparently this characteristic is shared across eukaryote organisms. Hettiarachchi and Saitou (2016) reported that conserved non-coding sequences (CNSs) in Diptera and vertebrates are poor in GC content. However, it is not clear the meaning of this skewed number of the bases in the genetic material. Tajima’s neutrality test values (D) for COI, ND4 and concatenated (COI-ND4) sequences were positive (3.60131, 3.67790, 2.11123, respectively) and significant (P<0.05) that suggests balancing selection or population substructure (McVean 2002). According to Weedall and Conway (2010) a positive Tajima’s D can either indicate balancing selection in the sample, or that sequences were sampled across divergent populations. The fixation index (F_ST_) of COI (0.12328), ND4 sequences (0.22688), and concatenated analysis of sequences of both genes (0.12531) indicate genetic differentiation of the *Ae. aegypti* populations (Wright 1978). The positive value of neutrality tests (Fu & Li, 1993) of COI sequences (F* = 2.62213, D*= 1.51245), ND4 sequences (F* = 2.4672, D*= 1.23977) and concatenated sequences of both genes (F* = 2.1071, D*= 1.64213), which may indicate balancing selection or population substructure. These positive values indicate a lack of singletons (mutations appearing only once among the sequences), which agree with the positive Tajima values, though this is not always necessarily the case (Fu & Li, 1993). However, the two tests differ in their sensitivity to the number of mutations of intermediate frequency, so that one test can be significant when the other one is not. This was observed in the values of D* (1.23977, not significant) of ND4 sequences of all population samples.

When comparing the three geographical regions (Galápagos, Amazon basin and Pacific coast) all the values of neutrality tests were positive, which may indicate balancing selection or population substructure. AMOVA indicate that there is not difference among the groups (geographical regions). The negative value is because variance components in AMOVA are actually defined as covariances, negative values can occur (Excoffier and Lischer 2010). The values obtained in the analysis by region (Pacific coast, Amazon basin and Galápagos) of ND4 showed population structuring (Costa-da-Silva et al. 2005).

The origin of the populations of *Aedes aegypti* in Ecuador show African genetic origin and widely present in several countries in the Americas. One of the genetic variants is more common in all the localities. The two haplotypes are distributed indistinctly in the three geographical sampled areas. A more detailed spatial and temporal sampling and more genes may be analyzed to reach conclusions about the populations of *Ae. aegypti* in Ecuador. The genetic identity of the mosquito populations may have a roll in vector competence transmitting arboviruses, including DENV, ZIKV, CHIKV and eventually YFV. The knowledge of the vector genetic variation may contribute to understand the epidemiology of arboviral diseases, routes of vector and viral dissemination and aid in the design of effective control strategies.

## Acknowledgments

We thank SENESCYT for grants PIC-12INH-002, PIC-12INH-003. We thank Andrea Fernández and Victoria Nipaz for the help processing the samples.

